# Dual Targeting of MAPK Signaling and Senescence-Associated Survival Pathways Overcomes Vincristine Resistance in Fusion-Negative Rhabdomyosarcoma

**DOI:** 10.1101/2025.07.29.667542

**Authors:** Yadong Wang, Andrea Largent, Eleanor Y. Chen

**Affiliations:** Department of Laboratory Medicine and Pathology, University of Washington, Seattle, WA

## Abstract

Rhabdomyosarcoma (RMS) is the most common pediatric soft tissue sarcoma, with limited therapeutic options for relapsed or metastatic disease. Fusion-negative (FN) RMS, the predominant subtype, frequently harbors mutations activating the RAS/MAPK pathway, yet the mechanisms underlying chemoresistance remain poorly defined. Here, we generated vincristine-resistant FN RMS cell lines through long-term dose escalation and identified increased MAPK signaling activity, evidenced by elevated phosphorylated ERK levels *in vitro* and in patient tumors post-chemotherapy. MEK inhibition via trametinib or MEK1 knockdown significantly impaired growth of resistant cells and enhanced vincristine efficacy in zebrafish tumor models, reducing tumor burden and relapse. Mechanistically, trametinib induced cell cycle arrest and senescence—marked by p21 upregulation—in resistant cells. This effect was dependent on suppression of MYC, as MYC disruption promoted senescence while MYC overexpression rescued the phenotype. Senescent cells remained partially viable via BCL-XL-mediated survival, rendering them sensitive to the BCL-XL inhibitor, A-1155463. Combined trametinib and A-1155463 significantly reduced growth of resistant RMS cells. Our findings highlight the MEK/MYC/p21 axis and BCL-XL dependency as key vulnerabilities in therapy-resistant FN RMS and support dual targeting of MAPK and senescence-survival pathways as a rational therapeutic strategy.

**STATEMENT OF SIGNIFICANCE:** Therapeutic resistance remains a major barrier to improving survival outcomes in fusion-negative rhabdomyosarcoma (FN RMS), the most common pediatric soft tissue sarcoma. This study identifies enhanced MAPK signaling and MYC-driven suppression of p21 as key mechanisms by which FN RMS cells evade vincristine-induced senescence. Moreover, we reveal that senescent-resistant cells rely on BCL-XL for survival, uncovering a new vulnerability that can be exploited therapeutically. Dual targeting of the MEK/MYC/p21 axis and BCL-XL– mediated survival pathways significantly suppresses growth of vincristine-resistant FN RMS cells. These findings provide a compelling rationale for combination therapies that simultaneously disrupt oncogenic signaling and senescence escape mechanisms in treatment-refractory FN RMS.

## INTRODUCTION

Rhabdomyosarcoma (RMS) is a malignant soft tissue tumor arising from impaired muscle differentiation and is the most common pediatric soft tissue sarcoma, with approximately 300 cases annually in the United States^1,2^. RMS is classified into two major molecular subtypes: fusion-negative (FN) RMS and fusion-positive (FP) RMS. FN RMS—the more prevalent subtype—is driven by mutations in the receptor tyrosine kinase/RAS/PIK3CA signaling axis in over 90% of cases^3^. In contrast, FP RMS is defined by a translocation involving PAX3 or PAX7 fused to FOXO1^4,5^.

Despite aggressive treatment with surgery, chemotherapy, and radiation, outcomes remain poor for patients with relapsed or metastatic disease, with 3-year survival rates under 30%^6^. Moreover, standard therapies cause significant long-term morbidity and secondary malignancies. No new effective treatments have emerged for relapsed RMS over the past three decades, underscoring the urgent need to elucidate the biological mechanisms underlying therapy resistance to develop more effective and less toxic treatment options.

Current understanding of chemotherapy resistance in RMS remains limited. Prior studies have shown that RMS cells treated with vincristine or doxorubicin retain a MYOD1-positive population^7^, which may contribute to persistence. MYOD1, a master regulator of skeletal muscle differentiation, is aberrantly regulated in RMS, leading to repression of myogenic genes and promotion of proliferation^8,9^. Other studies have identified elevated anti-apoptotic proteins and activation of survival pathways, such as GP130/STAT and Hedgehog, in therapy-resistant RMS^10,11^. These data suggest that therapy resistance is accompanied by dynamic shifts in transcriptional and signaling programs, but the cellular mechanisms enabling long-term survival of therapy-resistant RMS cells remain incompletely characterized.

We previously showed that vincristine treatment induces a primitive, stem-like phenotype in RMS cells^12^. Building on these findings, we now investigate the underlying biological mechanisms essential for maintaining the survival of therapy-resistant FN RMS cells. Our study identifies upregulated MAPK signaling as a key driver of senescence bypass in vincristine-resistant FN RMS. We further demonstrate that the MEK/MYC/p21 axis and dependence on BCL-XL-mediated survival represent targetable vulnerabilities. These findings suggest that co-targeting MAPK signaling and senescence-survival pathways is a promising therapeutic strategy for treatment-refractory FN RMS.

## METHODS/MATERIALS

### Generation of Vincristine-Resistant RMS Cell Lines

RD and SMS-CTR FN RMS cell lines were initially treated with vincristine at a dose (1-1.5 nM) that killed around 80-90% of cells. The remaining cells were maintained in long-term culture with weekly change of media (10% FBS and 1% penicillin/streptomycin in DMEM) until they grew to >50% confluency and then split to allow regrowth before restarting vincristine at a 10-20% increase in dosage. Incremental increases in drug dosage was continued over the course of 9 months to a year to reach an at least 100-fold increase in the IC50 of vincristine. Vincristine-resistant lines were maintained in 25-35 nM of vincristine prior to plating for assays. All cell lines were previously authenticated by STR profiling and tested for mycoplasma contamination.

### CRISPR/Cas9 and shRNA Gene Targeting in Human RMS Cell Lines

CRISPR/Cas9 targeting was accomplished by transducing RMS cells with lentivirus expressing safe-harbor control^13^ or gene-specific double guide RNAs (gRNAs) against *MYC*^12^ and Cas9. Gene-specific short hairpin RNAs (shRNAs) were obtained from Sigma (Table S1 – shRNA sequences). Cells transduced with lentivirus were plated for cell counts and cell-based assays following antibiotic selection 6 days post transduction. Cloning of Cas9 and gRNA expression vectors was performed as described previously ^13^.

### RNA Sequencing

RNA was extracted from parental and vincristine-resistant RD cells using the Qiagen RNeasy mini kit (catalog number: 74104) per manufacturer’s instructions. The RNA samples (3 biological replicates each) were submitted to the RNA sequencing service at Azenta Life Sciences (Seattle, WA) for whole transcriptome library preparation and sequencing.

### Cell Viability Assays

RMS cell growth was quantified by cell counting or the ATP-based CellTiter-Glo cell viability assay per manufacturer’s protocol (Promega, Madison, WI). Cells were plated in 96-well plates at 1000-2000 cells per well, and the luminescence was read using a microplate reader 1 day and 6 days post-plating (BioTek Synergy H1, Winooski, VT). Drug treatments were introduced 1 day post-plating.

### Immunohistochemistry (IHC) and Immunofluorescence (IF)

Archival paraffinized tissue blocks of human RMS tumor samples were retrieved using an approved IRB protocol (#9565) at University of Washington. De-identified tissue sections at 4-micron thickness were subjected to IHC. Briefly, following deparaffinization and antigen retrieval with the Tris-EDTA buffer for cleaved Caspase 3 antibody (Cell Signaling) slides and phosphate-buffered saline (PBS) for the phosphor-ERK antibody (Cell Signaling) slides for 15 minutes while boiling, the slides were stained with appropriate antibodies using standard IHC protocol. DAB (3,3-diaminobenzidine) HRP substrate was used for signal detection.

Immunofluorescent staining was performed as previously described^14^. The antibodies used for IHC and IF are detailed in table S1.

### Real-time Apoptosis Detection in Cultured RMS cells

For detecting apoptosis in cultured cells, adherent RMS cells or spheroids were incubated with NucView® 488 Caspase 3 substrate (Biotium, Fremont, CA), diluted to 2 μM in 500 μL medium, for 30 mins prior to imaging by microscopy.

### Western Blots

Human cell lysates were prepared in RIPA buffer with protease inhibitors (Santa Cruz Biotechnology, Dallas, TX) and 2x Laemmli buffer supplemented with β-mercaptoethanol (Hercules, CA), electrophoresed on a 4-15% gradient SDS-polyacrylamide gel (Bio-Rad, Hercules, CA), and transferred to a PVDF membrane using the TurboTrans-Blot (Bio-Rad, Hercules, CA). Blots were blocked in 5% milk (or 5% bovine serum albumin for the phospho-ERK antibody) in Tris-buffered saline with Tween 20 (TBST) and probed against corresponding antibodies as listed in table S1.

### Flow Cytometry for Cell Cycle Analysis

1 million human RMS cells were placed in Eppendorf tubes and resuspended in 100 microliters of PBS or FACS buffer [PBS (Gibco), 1% bovine serum albumin (BSA, Fisher Bioreagents), 0.5mM EDTA (Invitrogen, 0.5M)] containing flow antibodies for 20 minutes at 4°C. Cells were then fixed in a 4% paraformaldehyde solution (Thermo Fisher Scientific, Waltham, MA) for 20 minutes at 4°C before staining with DAPI (BD Biosciences) for 30 minutes at room temperature.

### **β**-Gal Staining

β-Gal staining was performed as previously described^15^. Imaging was performed using a BioTek Cytation 7 Cell Imaging Multimode Reader (Agilent, Santa Clara, CA).

### Quantitative RT-PCR (qPCR)

Human or zebrafish cells were lysed in TRIzol reagent (Thermo Fisher Scientific, Waltham, MA) and RNA was isolated per the manufacturer’s protocol. Approximately 1 microgram of RNA was used for cDNA synthesis using the High-Capacity cDNA Reverse Transcription kit (Thermo Fisher Scientific, Waltham, MA). SYBR Green-based quantitative PCR was subsequently performed using gene-specific primers (Table S1) in a CFX Connect Real-time PCR Detection System light cycler (Bio-Rad, Hercules, CA).

### Drug Treatment Studies in the Zebrafish RMS Model

Zebrafish were maintained in a shared facility at the University of Washington under protocol #4330-01 approved by the Institutional Animal Care and Use Committee at the University of Washington. Liquid nitrogen-preserved primary tumors with *rag2*:*KRASG12D* co-expressing mCherry were expanded by transplantation into 3-4 syngeneic adult CG1 fish. Tumor cells were then harvested by dissection and dissociation and transplanted subcutaneously into the peritoneal cavity at 25,000 cells per fish. A drug treatment study was performed on engrafted tumor-bearing tumor fish ∼10 days post-transplantation for 7 days. Drugs reconstituted in DMSO [vincristine (Sigma Aldrich) at 0.1 mg/kg and trametinib (10 mg/kg)] and vehicle (DMSO) were delivered on day 0 and day 4 in a 5-microliter volume using a Hamilton syringe needle via the intraperitoneal route. Imaging of tumor-bearing fish was performed on day 0 and day 7 using a Nikon fluorescent dissecting scope (Nikon Instruments Inc., Melville, NY). The tumor volume change was quantified using ImageJ software as previously described^14^.

## RESULTS

### Increased activity of MAPK pathway in vincristine-resistant FN RMS cells

To investigate the underlying biological features of vincristine-resistant FN RMS cells, we generated two therapy-resistant FN RMS lines (RD and SMS-CTR) through gradual vincristine dose escalation over 9–12 months, achieving at least a 100-fold increase in IC50 (see schematic in Fig. 1A; dose–response curves in Fig. S1A, B). Vincristine-resistant RD cells were subsequently collected for RNA sequencing. Pathway and oncogenic signature analyses revealed upregulation of MAPK signaling components, including RAS and RAF (Fig. 1B).

**Figure 1.**
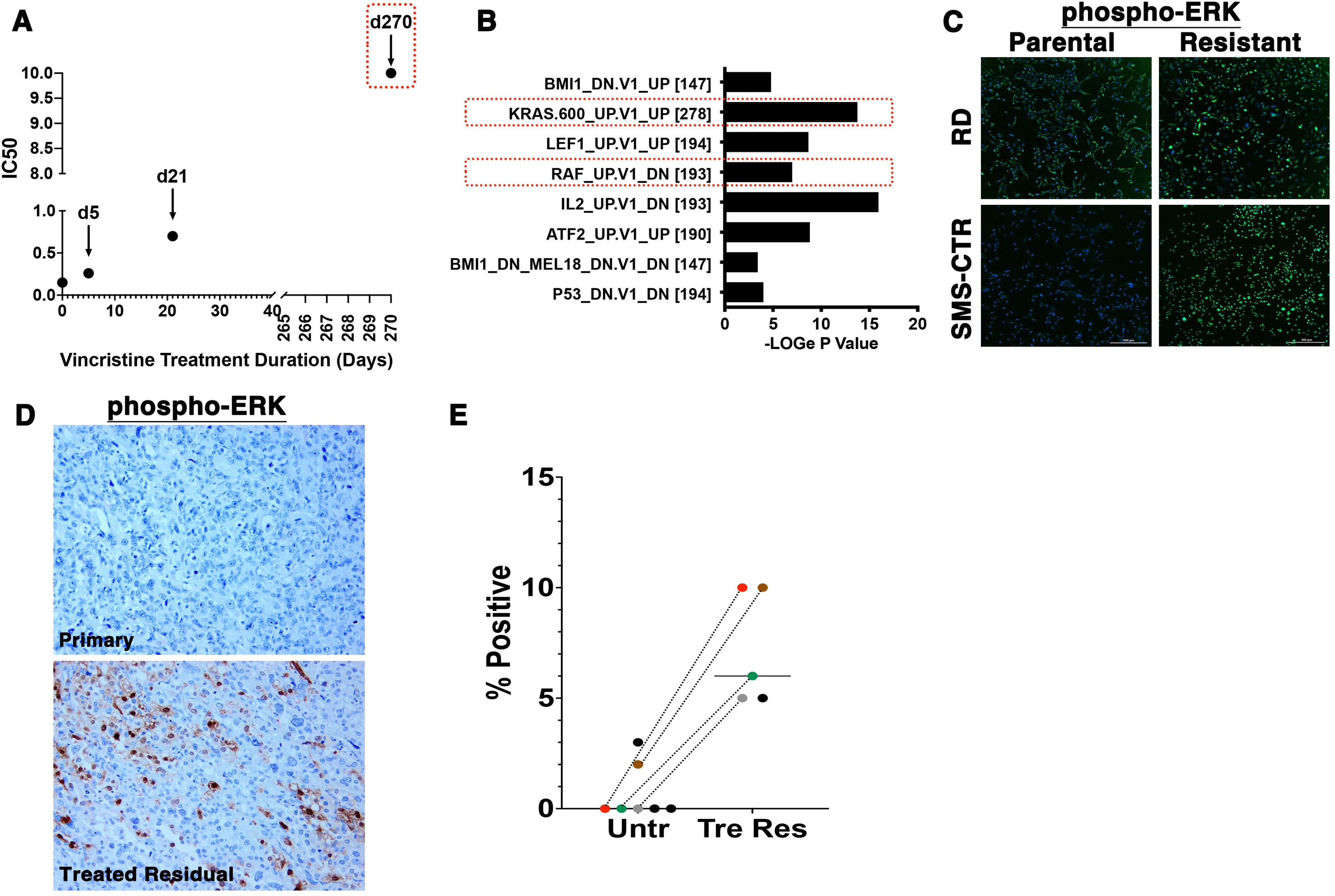
Increased MAPK signaling activity in therapy-resistant FN RMS cells. (A) Schematic of the generation of vincristine-resistant FNRMS cells in culture via gradual dose escalation over time. The Y-axis represents IC50 (nM) of RMS cells treated with vincristine; the X-axis represents duration of culturing in days. (B) Oncogenic signature analysis using the Broad Institute Molecular Signatures Database, based on gene expression profiles of vincristine-resistant RD cells from RNA sequencing. (C) Representative immunofluorescence (IF) images for phospho-ERK invincristine-sensitive parental (P) and resistant RD and SMS-CTR cells (Vinc R). Green = phospho-ERK, blue = DAPI. (D) Representative immunohistochemistry (IHC) images for phospho-ERK on formalin-fixed paraffin-embedded (FFPE) tissue sections from a FN RMS patient: primary tumor (top) and matched residual tumor post-chemotherapy (bottom). (E) Quantitative summary of phospho-ERK IHC in primary and post-chemotherapy FN RMS FFPE patient samples. Matched samples are color-coded and connected by dashed lines. Untr, untreated; Tre Res, treated residual.

To validate MAPK pathway activation, we performed immunofluorescence analysis, which showed elevated phosphorylated ERK (pERK) expression in both vincristine-resistant RD and SMS-CTR cells compared to parental lines (Fig. 1C). Similarly, in archival formalin-fixed, paraffin-embedded (FFPE) tumor samples from five FN RMS patients—including four with matched pre-treatment samples—residual post-chemotherapy tumors were enriched for pERK-positive cells (Fig. 1D, E). These findings suggest that MAPK signaling plays a key role in driving therapy resistance in FN RMS.

### Therapeutic efficacy of MEK inhibition as monotherapy and in combination with vincristine

Given the activation of MAPK signaling, we assessed whether MEK inhibition could suppress growth of vincristine-resistant FN RMS cells. Treatment of vincristine-resistant RD and SMS-CTR cells with trametinib, a MEK inhibitor, significantly reduced cell proliferation in a dose-dependent manner (Fig. 2A). Similarly, MEK1 knockdown using two independent shRNAs also impaired the growth of resistant RD and SMS-CTR cells (Fig. 2B, 2C). Notably, vincristine-resistant RD cells demonstrated greater sensitivity to MEK1 inhibition compared to parental cells, while this differential effect was not observed in SMS-CTR cells (Fig. 2B).

**Figure 2.**
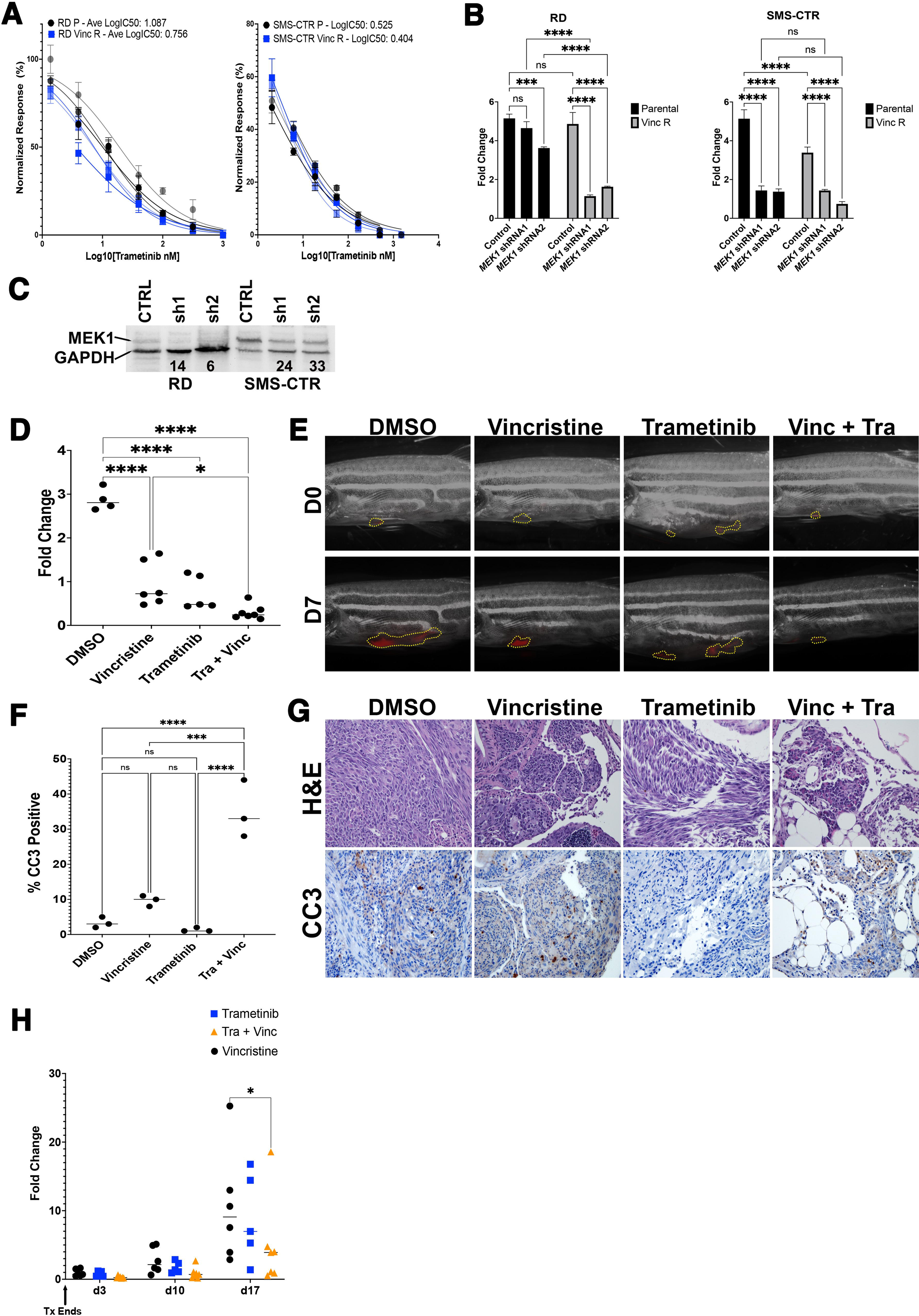
Trametinib reduces growth and relapse in vincristine-resistant FN RMS. (A) Dose-response curves of parental and vincristine-resistant RD and SMS-CTR cells treated with trametinib normalized to DMSO (vehicle control) cells. P, parental, Vinc R, vincristine-resistant. Each dose was tested in 3 replicate wells. Shown are results from one of at least three independent experiments. (B) Summary of CellTiter-Glo viability assays in RD and SMS-CTR cells (parental and vincristine-resistant) transduced with two independent MEK1-targeting shRNAs. Scrambled shRNA was used as a control. Relative fold change in luminescence over 5 days was calculated, with treatment starting 6 days post-transduction. (C) Western blot analysis of MEK1 protein levels at 5 days post-shRNA transduction. GAPDH served as loading control. Normalized band intensity for MEK is indicated below each lane. (D) Quantification of relative tumor volume changes from zebrafish drug treatment studies. Each dot represents one animal. One-way ANOVA with multiple comparisons correction; ns, not significant; *P < 0.05; ****P < 0.0001. (E) Representative images of zebrafish engrafted with KRAS(G12D)-induced FN RMS tumors treated with DMSO (vehicle), vincristine (0.1 mg/kg), trametinib (10 mg/kg), or the combination. Tumor-bearing zebrafish received intraperitoneal injections on days 0 and 4 over a 7-day period. (F) Quantification of cleaved Caspase-3 (CC3) IHC. The percentage of CC3-positive cells per whole tumor cross-section was measured using QuPath (n = 3 per condition). Statistical test used? ns, not significant; ***P < 0.001; ****P < 0.0001. (G) Representative tumor sections stained with H&E (top) and CC3 IHC (bottom). (H) Tumor volume measurements of relapsed tumors over time post-treatment. Two-way ANOVA with multiple comparisons correction; *P < 0.05.

To evaluate whether MEK inhibition enhances vincristine efficacy *in vivo*, we used a KRAS(G12D)-induced zebrafish FN RMS model. Primary RMS cells expressing the *mylz*:mCherry reporter were transplanted into syngeneic CG1 zebrafish, followed by treatment with vehicle (DMSO), trametinib (10 mg/kg), vincristine (0.1 mg/kg), or both drugs in combination for 7 days. Drug doses were titrated to ∼IC50 in vivo. Combination treatment significantly reduced tumor growth compared to either monotherapy (n = 4–7 per group, Fig. 2D, E). Apoptosis, assessed via cleaved Caspase-3 immunohistochemistry (IHC), was significantly increased in the combination group compared to either single-agent arm (Fig. 2F, G), supporting enhanced tumor cell death.

To test whether this regimen also prevented relapse, we monitored tumor regrowth post-treatment. By day 17, zebrafish receiving combined vincristine and trametinib showed significantly lower relapse frequency and slower growth of recurrent tumors than those treated with either drug alone (p < 0.05, Fig. 2H). These results indicate that dual targeting of MEK and vincristine may improve treatment durability in FN RMS.

### Trametinib treatment alters cell cycle progression and induces senescence in FN RMS cells

To investigate the cellular mechanisms underlying growth inhibition following MEK1 knockdown or trametinib treatment, we assessed alterations in cell cycle, apoptosis, and myogenic differentiation. DAPI-based flow cytometry revealed that vincristine-resistant RD and SMS-CTR cells exhibited a prolonged G0/G1 phase compared to parental cells (Fig. 3A). Trametinib treatment further exacerbated G0/G1 arrest in both parental and vincristine-resistant cells (Fig. 3A, B).

**Figure 3.**
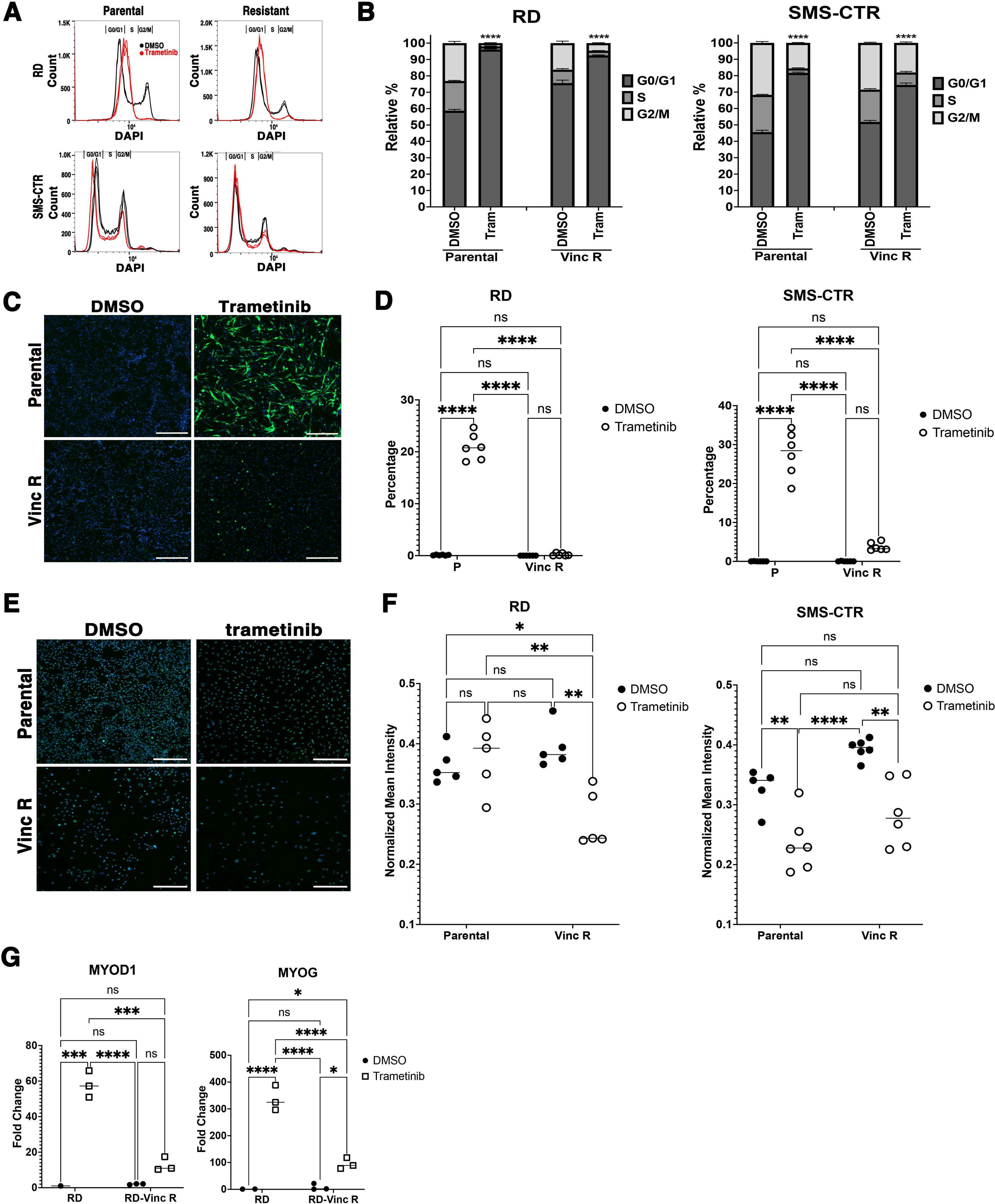
Trametinib induces cell cycle arrest in vincristine-resistant FN RMS cells. (A) Representative cell cycle histograms by DAPI-based flow cytometry. Black: DMSO; Red: trametinib (100 nM). (B) Summary of cell cycle distribution. (C) Representative IF images for RD cells stained with MF20 antibody, day 3 of drug treatment. Green = MF20, blue = DAPI. (D) Quantification of MF20 IF in RD and SMS-CTR cells (parental vs. vincristine-resistant). (E) Representative IF images for RD cells stained with the antibody for MYOD1, day 3 of drug treatment. (F) Quantification of MYOD1 IF in RD and SMS-CTR cells. (G) Quantitative RT-PCR of MYOD1 and MYOG, 24 hours of drug treatment. Error bars in (B, D, F, G) indicate standard deviation of 3 independent replicates. ns, not significant; *P < 0.05; ***P < 0.001; ****P < 0.0001 by two-way ANOVA with multiple comparisons correction.

We next evaluated apoptosis using live-cell imaging with NucView® 488 Caspase-3 substrate following 6 days after shRNA transduction or 3 days of trametinib treatment. Trametinib did not significantly increase apoptosis in either parental or vincristine-resistant RD or SMS-CTR cells compared to DMSO controls (Fig. S2 A, B). While MEK1 shRNA induced a statistically significant increase in apoptotic cells in vincristine-resistant RD cells compared to parental cells, the effect was modest (mean 1.2% vs. 0.3%, Fig. S2 C, D), suggesting that apoptosis contributes minimally to reduced cell viability.

Trametinib has previously been shown to induce terminal differentiation in FN RMS cells^16^. In agreement with prior findings, trametinib-treated parental RD and SMS-CTR cells exhibited increased MF20-positive differentiated muscle cells (Fig. 3C, D). However, trametinib-treated vincristine-resistant RD and SMS-CTR cells failed to undergo terminal differentiation to the same extent, as shown by reduced MF20 staining (Fig. 3C, D). Additionally, MYOD1 protein expression was decreased in resistant cells following trametinib treatment (Fig. 3E, F), and quantitative RT-PCR confirmed attenuated induction of MYOD1 and MYOG (20–30 fold lower) compared to parental cells (Fig. 3G).

Vincristine is known to induce senescence in various cancers through therapy-induced senescence (TIS) mechanisms^17,18^. Consistent with this, vincristine-resistant RD and SMS-CTR cells displayed enlarged, flattened morphology consistent with senescence (Fig. 4A). We used senescence-associated β-galactosidase (SA-β-gal) staining, a common marker of senescence, and found that vincristine-resistant RD and SMS-CTR cells had more β-gal-positive cells than treatment-naïve parental cells (Fig. 4B, C, S3A).

**Figure 4.**
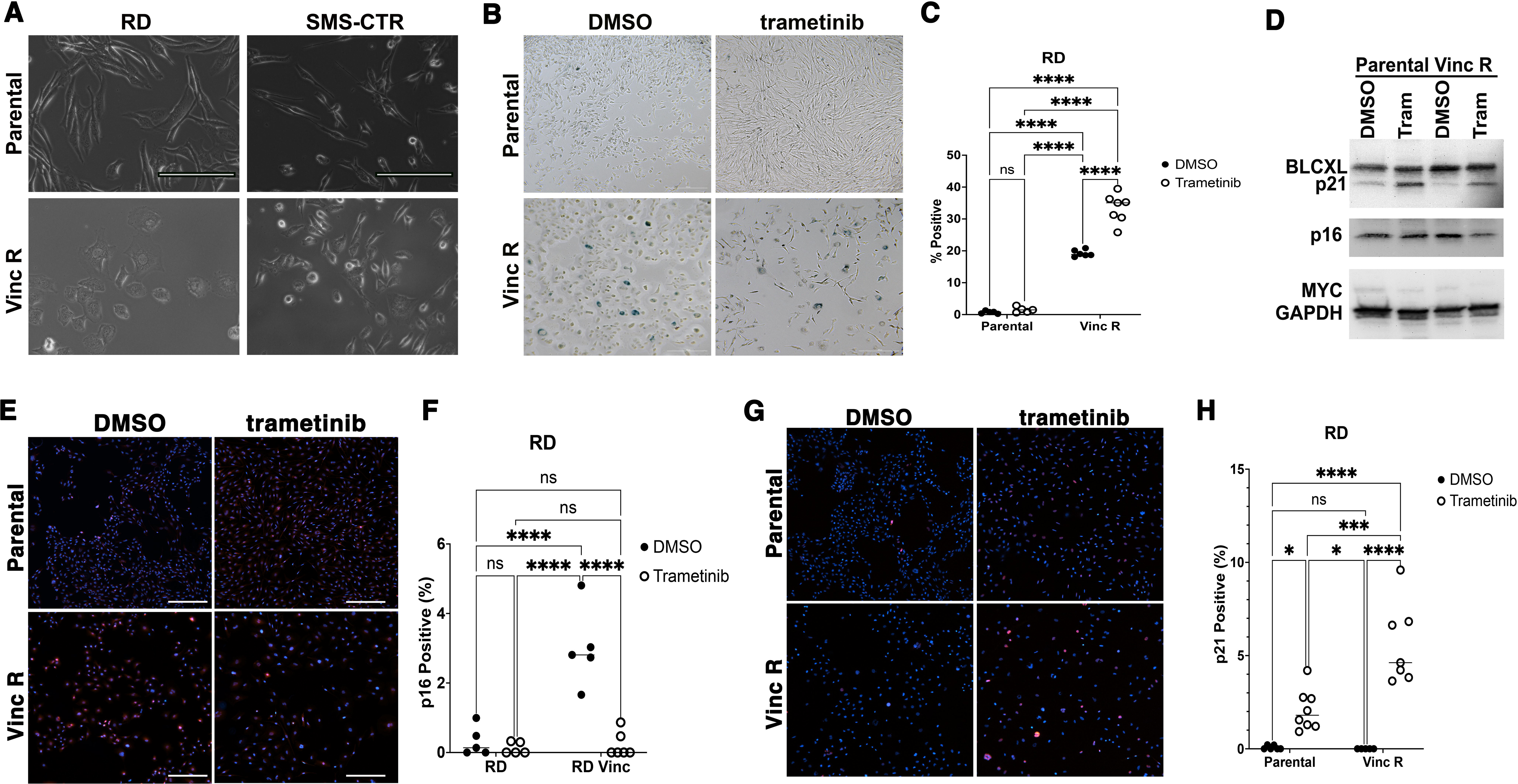
Trametinib induces senescence in vincristine-resistant FN RMS cells. (A) Bright-field images of parental (top) and vincristine-resistant (bottom) RD and SMS-CTR cells. Scale bar = 200 μm. Vincristine-resistant cells display larger, irregular morphology. (B) β-Gal staining of RD parental and resistant cells after 3 days of treatment with DMSO or trametinib (100 nM). (C) Quantification of β-Gal staining in RD cells (≥3 fields per condition at 100×). (D) Western blots of total lysates from RD? parental and resistant cells treated with DMSO or trametinib, 24 hours of drug treatment. GAPDH = loading control. (E) IF for p16 in RD cells, 72 hours of drug treatment. Red = p16, blue = DAPI. (F) Quantification of p16 IF in RD cells (≥3 fields per condition at 10×). Average of intensity per cell? ns, not significant; ****P < 0.0001 by two-way ANOVA with multiple comparisons correction. (G) IF for p21 in RD cells, 72 hours of drug treatment. Red = p21, blue = DAPI. (H) Quantification of p21 IF in RD cells (≥3 fields per condition at 10×). ns, not significant; *P < 0.05; ***P < 0.001; ****P < 0.0001 by two-way ANOVA with multiple comparisons correction.

Given that trametinib altered the cell cycle without inducing apoptosis or robust differentiation, we then asked whether it also promoted senescence in vincristine-resistant cells. Indeed, trametinib-treated vincristine-resistant RD and SMS-CTR cells showed a significant increase in β-gal-positive senescent cells compared to DMSO-treated controls (Fig. 4B, C, S3A). Parental cells did not exhibit this increase, supporting a resistance-associated senescence phenotype.

We next asked whether p21 or p16 contributed to this senescence-associated cell cycle arrest. Western blot and immunofluorescence analysis showed that trametinib-treated vincristine-resistant RD and SMS-CTR cells had increased p21 expression but reduced p16 levels (Fig. 4D– H, Fig. S3B-D). These findings suggest that p21, rather than p16, is the primary driver of trametinib-induced senescence in vincristine-resistant FN RMS cells.

### MYC represses p21 and suppresses trametinib-induced senescence in FN RMS

MYC is a known downstream effector of MAPK signaling and a critical regulator of proliferation. We previously showed that MYC promotes survival of vincristine-resistant FN RMS cells. Based on our observation that trametinib induces senescence and increases p21, we hypothesized that MYC may suppress p21 expression to block senescence.

We first confirmed that trametinib treatment reduced MYC protein levels in RD and SMS-CTR cells by Western blot (Fig. 4D). Next, we disrupted MYC using lentiviral CRISPR-Cas9 with validated dual gRNA constructs, which we have shown previously to effectively deplete MYC protein^12^. MYC disruption led to upregulation of CDKN1A (p21) mRNA, while CDKN2A (p16) levels remained unchanged (Fig. 5A), supporting a specific MYC–p21 axis.

**Figure 5.**
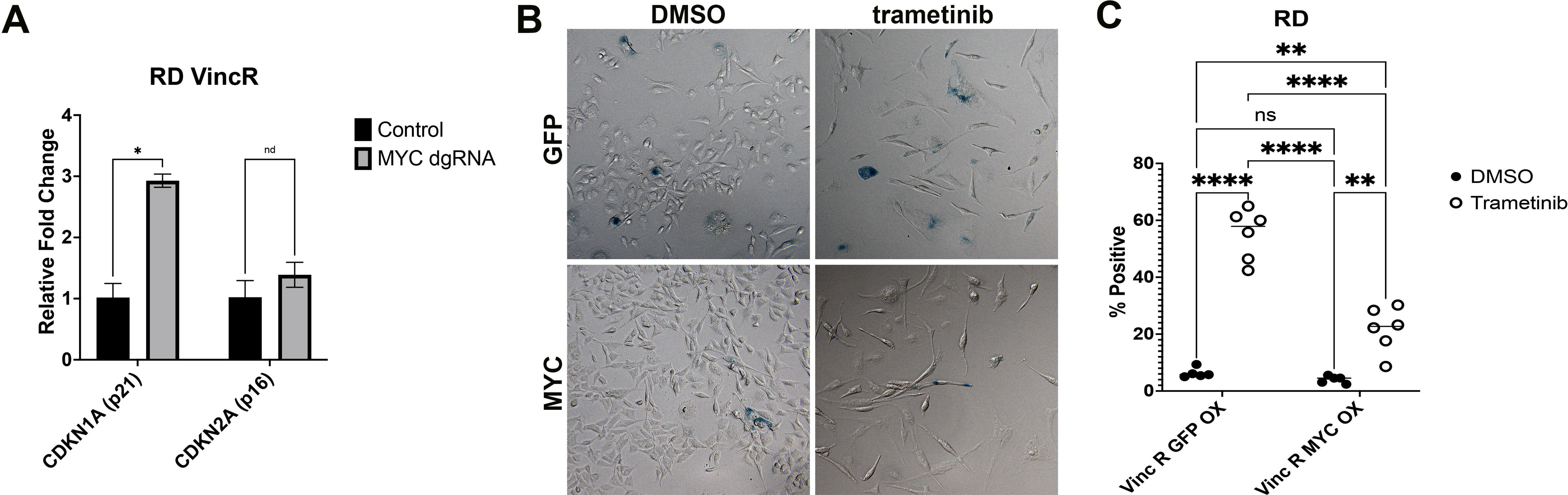
MYC alleviates trametinib-induced senescence in vincristine-resistant RMS cells. (A) qRT-PCR in vincristine-resistant RD cells following MYC disruption using Cas9 and dual gRNAs. RNA was collected 6 days post-transduction. Error bars = standard deviation. ns, not significant; *P < 0.05 by unpaired t-test. (B) β-Gal staining of vincristine-resistant RD cells transduced with GFP control or MYC-overexpression lentivirus and treated with DMSO or trametinib (100 nM) for 3 days. (C) Quantification of β-Gal staining (6 fields per condition at 20×). **P < 0.01; ****P < 0.0001 by two-way ANOVA with multiple comparisons correction.

To test whether MYC regulates senescence in this context, we overexpressed MYC in RD cells and found that MYC overexpression alleviated trametinib-induced senescence, as measured by reduced β-gal-positive cell frequency (Fig. 5B, C). These findings demonstrate that MYC suppresses p21-mediated senescence and is required for resistance to trametinib-induced cell cycle arrest in FN RMS.

### BCL-XL supports survival of senescent, vincristine-resistant FN RMS cells and represents a targetable vulnerability

While a subset of vincristine-resistant cells exhibited senescence markers—such as altered morphology and increased β-galactosidase staining (Fig. 4A, B)—the continued proliferation of the population suggested that some cells escape or bypass senescence-associated arrest. Prior studies in colorectal cancer have shown that escape from p21-mediated senescence can result in dependence on anti-apoptotic proteins, including BCL-XL and MCL1^19^.

To test whether vincristine-resistant FN RMS cells develop dependence on BCL-XL, we first examined protein expression following trametinib treatment. Western blot analysis revealed persistent BCL-XL expression in vincristine-resistant RD and SMS-CTR cells after trametinib exposure (Fig. 4D). Importantly, these resistant cells were more sensitive to the selective BCL-XL inhibitor A-1155463 than parental cells, as demonstrated by dose-response assays (Fig. 6A, B).

**Figure 6.**
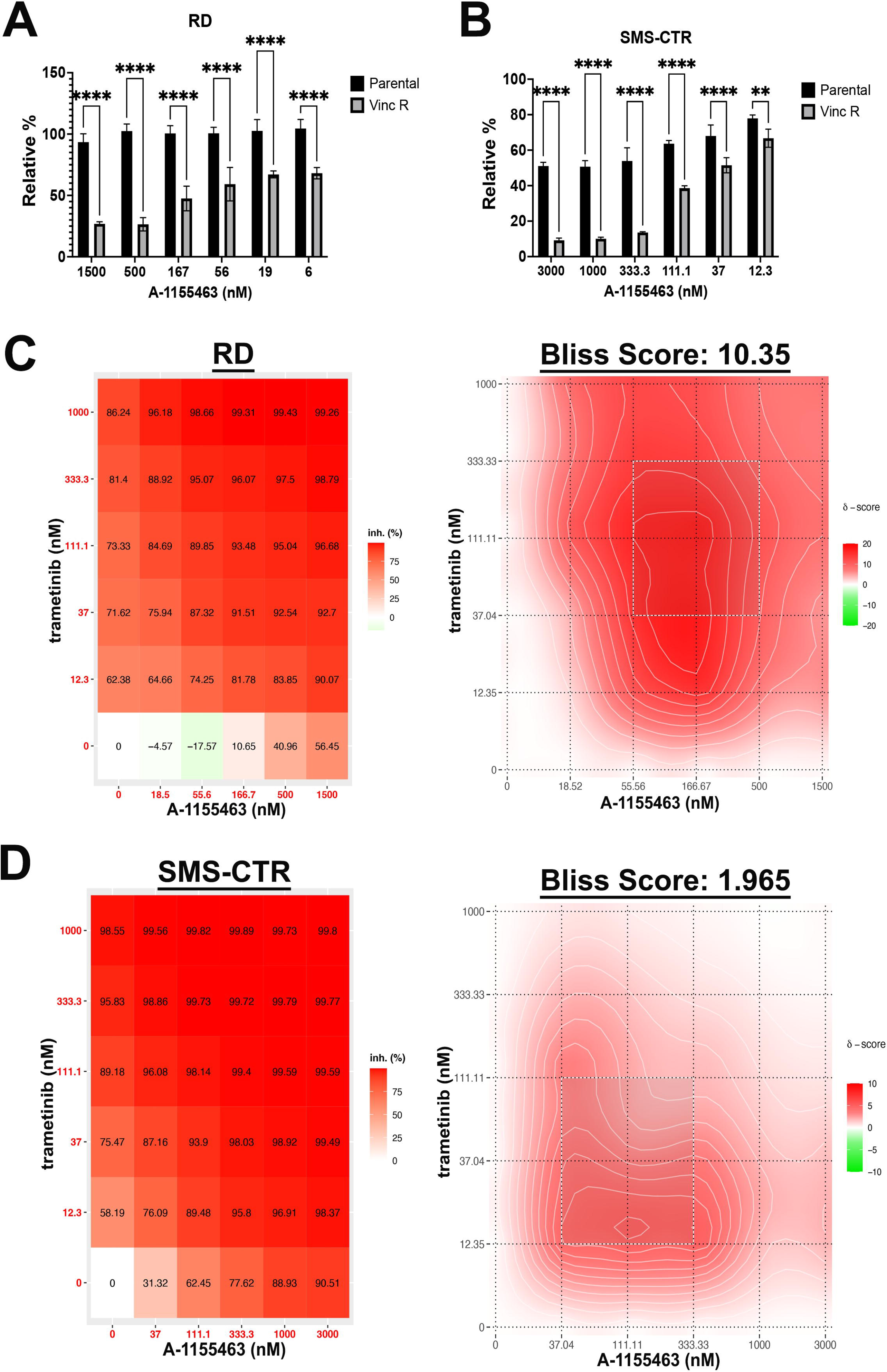
Combined MEK and BCL-XL inhibition reduces growth of vincristine-resistant FN RMS cells. (A, B) CellTiter-Glo assays assessing trametinib and A-1155463 dose responses in parental and vincristine-resistant RD (A) and SMS-CTR (B) cells. Luminescence normalized to DMSO controls. (C, D) Heatmaps (left) and 2D synergy contour plots (right) from 5-day combination treatment assays using trametinib and A-1155463. Bliss synergy scores and plots were generated using SynergyFinder (www.synergyfinder.org).

Given the dual dependencies on MEK signaling and BCL-XL-mediated survival, we hypothesized that combination targeting of these pathways would enhance therapeutic efficacy. Indeed, combination treatment with trametinib and A-1155463 at multiple dose combinations significantly inhibited growth of vincristine-resistant RD and SMS-CTR cells compared to single-agent treatment (Fig. 6C, D).

These findings indicate that senescence-associated survival mechanisms mediated by BCL-XL can be therapeutically exploited in vincristine-resistant FN RMS. Dual inhibition of MEK and BCL-XL represents a rational strategy to eradicate treatment-refractory tumor cells.

## DISCUSSION

In normal human fibroblasts, constitutive activation of MEK induces both p53 and p16 and is required for RAS-induced senescence of non-neoplastic human fibroblasts^20–22^. Oncogene-induced senescence resulting in cell cycle arrest is also a well-recognized phenomenon in cancer^23^. Conversely, inhibition of oncogenic MAPK signaling has also been shown to induce senescence in KRAS-mutant lung cancer, pancreatic cancer, and pilocytic astrocytoma^24–26^. The interplay between dose modulation of the driver oncogenic signaling pathway and the senescent response of cancer cells in the context of therapy resistance remains under characterized. In this study, we demonstrated that vincristine-resistant FN RMS cells and post-treatment persistent cells in a subset of FN RMS patient tumors exhibited enhanced MAPK signaling activity. While treatment-naïve parental cells underwent robust myogenic differentiation following trametinib treatment^16^, vincristine-resistant FN RMS cells did not show significant myogenic differentiation but instead underwent senescence-associated cell cycle arrest. Moreover, our findings, along with previously published data, suggest that the induction of a senescent state may be oncogene dosage–dependent. While excessive oncogene signaling can induce senescence, oncogene depletion can also conversely trigger senescence in neoplastic cells and may be therapeutically exploited to inhibit tumor growth.

We further demonstrated that senescence-associated cell cycle arrest in FN RMS was characterized by decreased MYC activity and increased expression of the cell cycle inhibitor p21. MYC was found to repress p21 expression, and overexpression of MYC abrogated trametinib-induced senescence. These findings underscore the pivotal role of MYC in sustaining proliferation of vincristine-resistant FN RMS cells by suppressing senescence-associated cell cycle inhibitors. Notably, prior work in early oral carcinogenesis revealed that MYC degradation is essential for triggering senescence as a protective mechanism against malignant transformation; in contrast, later-stage premalignant lesions (dysplasias) exhibit elevated MYC levels, suggesting that bypassing MYC degradation may be a prerequisite for senescence escape and tumor initiation^27^. Our data suggest a similar paradigm in FN RMS, wherein MYC promotes continued proliferation in vincristine-resistant cells by facilitating evasion of therapy-induced senescence. Collectively, these findings support a model in which MYC functions as a key suppressor of oncogene-induced senescence, thereby promoting tumor initiation and progression.

While trametinib-induced senescence and cell cycle arrest offer promising therapeutic avenues for targeting vincristine-resistant fusion-negative (FN) rhabdomyosarcoma (RMS) cells, our study revealed that these resistant cells exhibit increased dependence on the anti-apoptotic protein, BCL-XL, compared to their treatment-naïve parental counterparts. BCL-XL has previously been shown to antagonize p21 and promote escape from senescence, thereby enhancing cancer cell survival^19^. These findings suggest that upregulation of anti-apoptotic pathways may contribute to resistance against trametinib in RMS. Notably, we demonstrated that combined treatment with trametinib and the selective BCL-XL inhibitor A-1155463 significantly suppressed tumor growth in vincristine-resistant FN RMS cells, outperforming either agent alone. Together, these results support a therapeutic strategy that co-targets MAPK signaling and anti-apoptotic mechanisms as a potentially effective approach for overcoming resistance in treatment-refractory FN RMS.

Our study has several limitations. Standard-of-care chemotherapy regimens for rhabdomyosarcoma typically involve multi-agent combinations, including vincristine, actinomycin and cyclophosphamide. However, our investigation focused exclusively on resistance mechanisms following prolonged vincristine exposure and did not account for potential interactions among agents in combination therapy. While this represents a limitation, it also offers a strength—by isolating vincristine, we were able to eliminate confounding effects from other drugs and dissect vincristine-specific resistance pathways. Additionally, it remains unclear whether fusion-positive (FP) RMS shares similar resistance mechanisms to vincristine as fusion-negative (FN) RMS, given their distinct genetic drivers. Future studies are warranted to evaluate whether other chemotherapeutic agents induce distinct resistance phenotypes and to determine whether FP and FN RMS exhibit divergent mechanisms of therapy resistance.

In summary, we identified the MEK/p21/MYC axis for promoting growth and increased dependence on anti-apoptotic pathways in vincristine-resistant FN RMS cells, likely linked to senescence bypass or escape. Thus, co-targeting the MAPK pathway and anti-apoptotic programs may offer an effective strategy for treating refractory RMS.

## ACKNOWLEDGMENT

The study was supported by Department of Defense Rare Cancers Research Program Idea Development Award (RA210198).

