## Supplementary figures and images for "Dual Targeting of MAPK Signaling and Senescence-Associated Survival Pathways Overcomes Vincristine Resistance in Fusion-Negative Rhabdomyosarcoma"

### Figure S1

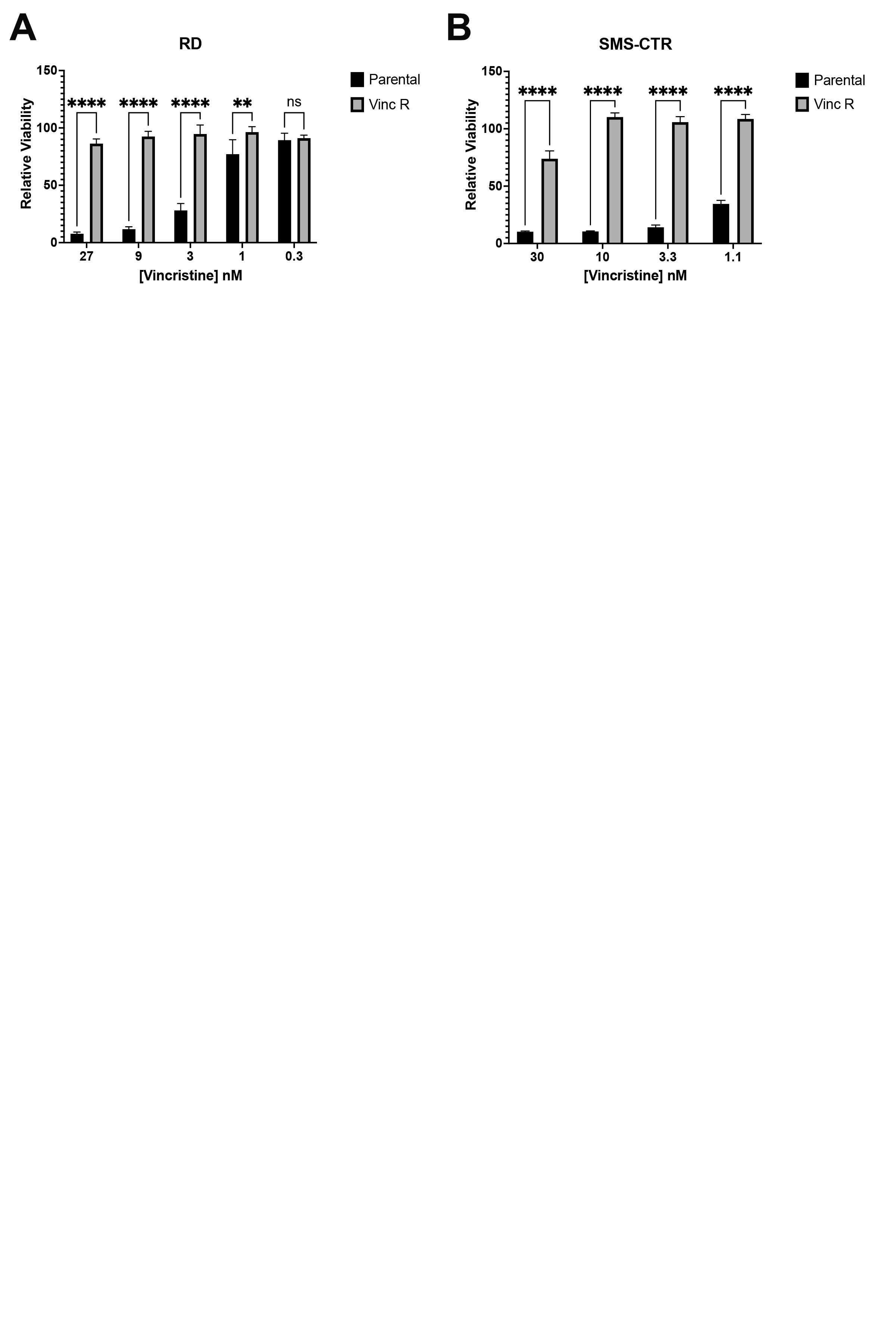

### Figure S2

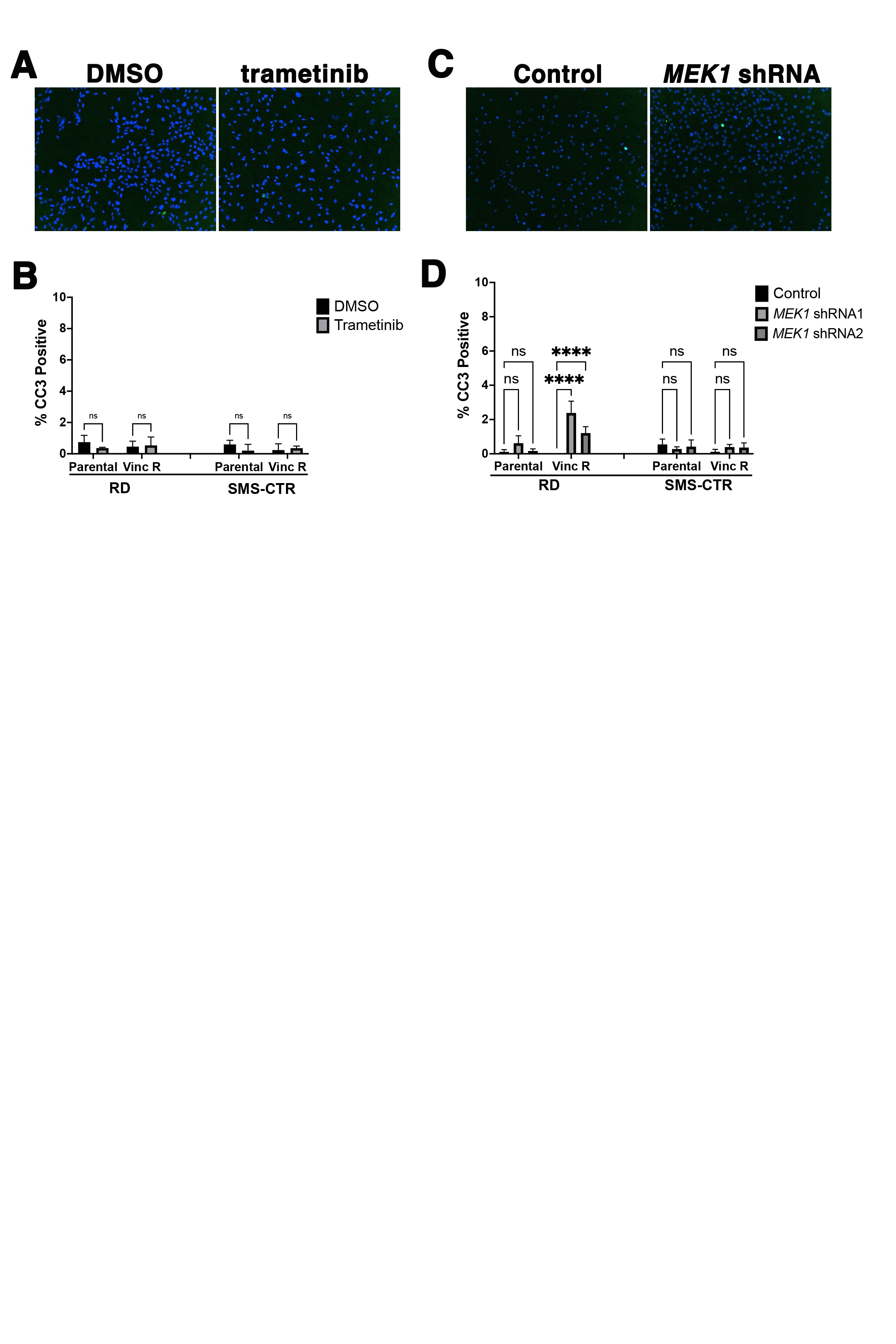

### Figure S3

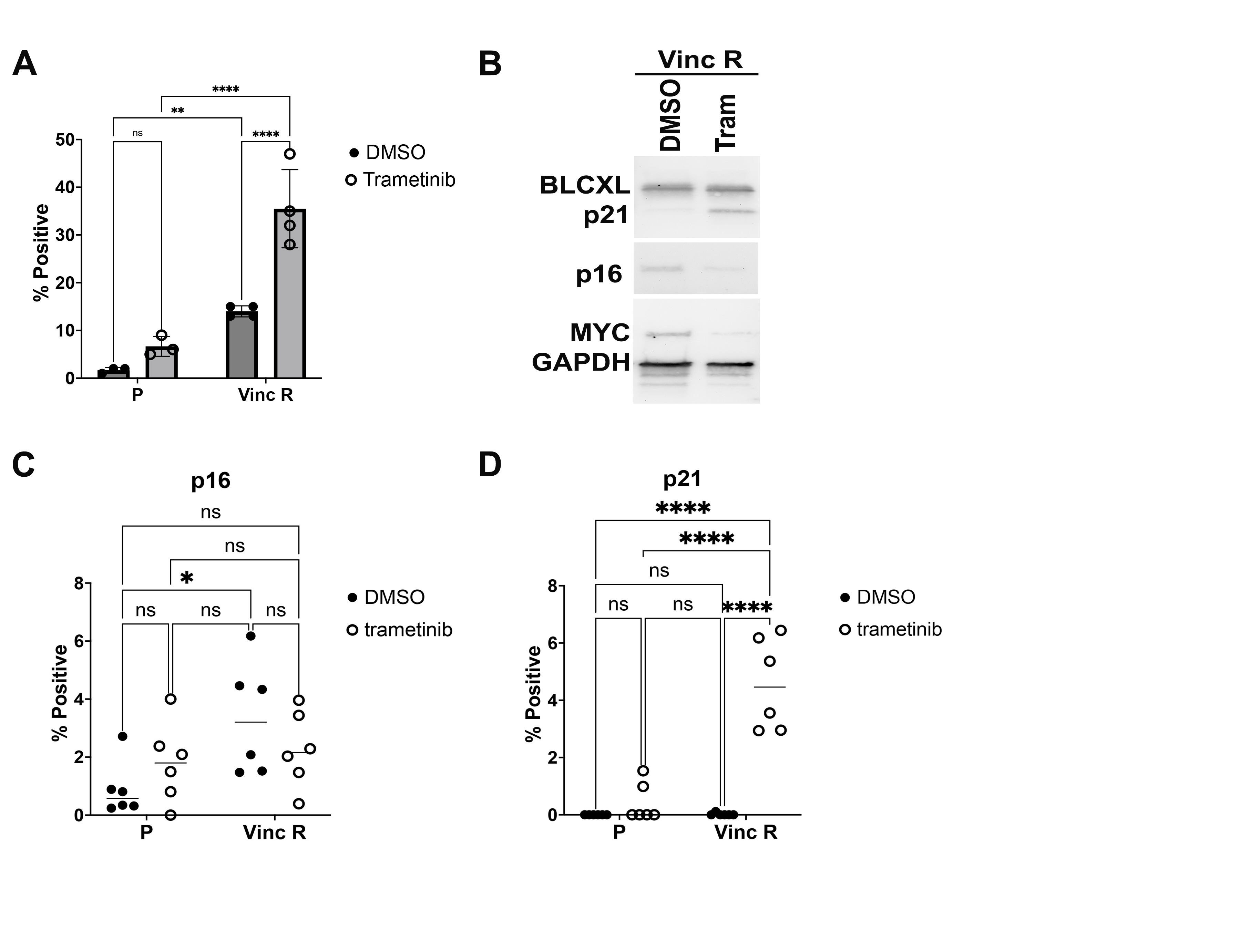
