## Supplementary material for "Dual Targeting of MAPK Signaling and Senescence-Associated Survival Pathways Overcomes Vincristine Resistance in Fusion-Negative Rhabdomyosarcoma": Table S1

| **Table S1. Antibodies, shRNAs and primers** |  |  |  |  |  |  |
| --- | --- | --- | --- | --- | --- | --- |
| **Antibodies for Immunofluorescence, Immunohistochemistry, Flow Cytometry and Western blots** | | | | | | |
| Protein | Source | Clone | Dilution | Company | Cat No | Assay |
| Cleaved Caspase-3 (Asp175) | Rabbit polyclonal |  | 1:300 | Cell Signaling | 9661 | IHC |
| Phospho-p44/42 MAPK (Erk1/2) (Thr202/Tyr204) | Rabbit monoclonal | 20G11 | 1:200 | Cell Signaling | 4370 | IHC |
| Phospho-p44/42 MAPK (Erk1/2) (Thr202/Tyr204) | Rabbit monoclonal | D13.14.4E | 1:300 | Cell Signaling | 9101 | IF |
| MYH1E (MF20) | Mouse monoclonal |  | 1:100 | DSHB | MF 20 | IF |
| p16-INK4A | Rabbit polyclonal |  | 1:500 | Proteintech | 10883-1-AP | IF |
| p16-INK4A | Rabbit polyclonal |  | 1:1000 | Proteintech | 10883-1-AP | WB |
| p21 Waf1/Cip1 | Rabbit monoclonal | 12D1 | 1:200 | Cell Signaling | 2947T | IF |
| p21 Waf1/Cip1 | Rabbit monoclonal | 12D1 | 1:1000 | Cell Signaling | 2947T | WB |
| MEK1 | Rabbit monoclonal | D2R1O | 1:1000 | Cell Signaling | 12671T | WB |
| BCL-xL | Rabbit monoclonal | 54H6 | 1:1000 | Cell Signaling | 2764T | WB |
| MYC | Rabbit monoclonal | E5Q6W | 1:1000 | Cell Signaling | 18583S | WB |
| GAPDH | Rabbit monoclonal | 14C10 | 1:2000 | Cell Signaling | 2118S | WB |
| **shRNAs** | | | | | | |
| Gene |  | Sequence |  |  |  |  |
| MEK1 | shRNA 1 | GCTTCTATGGTGCGTTCTACA | | |  |  |
| MEK1 | shRNA 2 | GAGGGAGAAGCACAAGATCAT | | |  |  |
| **RT-PCR Primers** | | | | | | |
| MYOD1 | Forward | AGCACTACAGCGGCGACT | | |  |  |
|  | Reverse | GCGACTCAGAAGGCACGTG | | |  |  |
| MYOG | Forward | CCTGCCGTGGGCGTGTAAGG | | |  |  |
|  | Reverse | GGACTGCAGGAGGCGCTGTG | | |  |  |
| CDKN2A (p16) | Forward | GTGGACCTGGCTGAGGAG | | |  |  |
|  | Reverse | CTTTCAATCGGGGATGTCTG | | |  |  |
| CDKN1A (p21) | Forward | CTGGGGATGTCCGTCAGAAC | | |  |  |
|  | Reverse | GTGACAGGTCCACATGGTCT | | |  |  |
