## Supplemental Figure Legends for "Dual Targeting of MAPK Signaling and Senescence-Associated Survival Pathways Overcomes Vincristine Resistance in Fusion-Negative Rhabdomyosarcoma"

**Figure S1. Vincristine-resistant FN RMS cells show reduced sensitivity to vincristine compared to parental cells.** (A, B) Dose-response curves comparing sensitivity of parental cells and vincristine-resistant cells in RD (A) and SMS-CTR (B) lines. **P < 0.01; ****P < 0.0001 by unpaired t-test, ns: not significant.

**Figure S2. Apoptosis is not a significant contributor to reduced cell growth secondary to *MEK1* shRNA or trametinib.** Apoptosis analysis via live imaging using with NucView® 488 Caspase-3 substrate was performed 6 days after shRNA transduction or 5 days post-trametinib (100 nM) treatment. (A,C) representative RD cell images. (B,D): quantification from three fields per condition in triplicate wells. Error bars in B and D indicate standard deviation of 3 independent replicates. ns, not significant; *P < 0.05; ****P < 0.0001 by two-way ANOVA with **multiple comparisons correction.**

**Figure S3. Trametinib exacerbated senescence in vincristine-resistant SMS-CTR cells via inductin of p21 expression.** (A) β-Gal staining of SMS-CTR parental and resistant cells after 5 days of treatment with DMSO or trametinib (100 nM). Quantification of β-Gal staining in SMS-CTR cells (≥3 fields per condition at 100×). (B) Western blots of total lysates from parental and resistant cells treated with DMSO or trametinib, 24 hours of drug treatment. GAPDH = loading control. (C) Quantification of p16 IF (≥3 fields per condition at 100×). (D) Quantification of p21 IF (≥3 fields per condition at 100×). ns, not significant; *P < 0.05; ****P < 0.0001 by two-way ANOVA with **multiple comparisons correction.**
